# Estimating *K* in Genetic Mixture Models

**DOI:** 10.1101/022988

**Authors:** Robert Verity, Richard A. Nichols

## Abstract

A key quantity in the analysis of structured populations is the parameter *K*, which describes the number of subpopulations that make up the total population. Inference of *K* ideally proceeds via the *model evidence*, which is equivalent to the likelihood of the model. However, the evidence in favour of a particular value of *K* cannot usually be computed exactly, and instead programs such as Structure make use of simple heuristic estimators to approximate this quantity. We show – using simulated data sets small enough that the true evidence can be computed exactly – that these simple heuristics often fail to estimate the true evidence, and that this can lead to incorrect conclusions about *K.* Our proposed solution is to use thermodynamic integration (TI) to estimate the model evidence. After outlining the TI methodology we demonstrate the effectiveness of this approach using a range of simulated data sets. We find that TI can be used to obtain estimates of the model evidence that are orders of magnitude more accurate and precise than those based on simple heuristics. Furthermore, estimates of *K* based on these values are found to be more reliable than those based on a suite of model comparison statistics. Our solution is implemented for models both with and without admixture in the software TrueK.

---

The detection and characterisation of population structure is one of the cornerstones of modern population genetics. Ever since Wright (1949) and his contemporaries (Malécot 1948) it has been recognised that genetic samples obtained from a large population may be better understood as a series of draws from multiple partially isolated subpopulations, or demes. While traditional methods (such as those based on the fixation index, *F*_*ST*_) assume that the allocation of individuals to demes is known *a priori*, many modern programs such as Structure (Pritchard *et al.* 2000; Falush *et al.* 2003a, 2007; Hubisz *et al.* 2009) take a different approach; attempting to infer the group allocation from the observed data. What makes this possible is the simple genetic mixture modelling framework used by Structure, together with the efficiency of Markov Chain Monte Carlo (MCMC) methods for sampling from this broad class of models.

However, even within the flexible framework of Bayesian mixture models, the number of demes (denoted *K*) is difficult to ascertain. While the allocation of individuals to demes is a parameter *within* a particular model, the value of *K* is fixed for a given mixture model, and so the problem of estimating *K* involves a comparison *between* models. One of the most common ways of comparing between models in a Bayesian setting is through the model evidence, defined as the probability of the observed data under the model. Often this quantity is used in a relative sense, with the ratio of evidence between competing models (the Bayes factor) being used to guide the choice of one model over another (Kass and Raftery 1995). However, computing the model evidence in complex, multi-dimensional models is not straightforward, and for this reason it is common to resort to simple heuristic estimators of the true evidence. These heuristics tend to have some direct mathematical connection to the model evidence, but also make certain simplifying assumptions in their derivation.

For example, in the original paper on which Structure is based, Pritchard *et al.* (2000) comment on the difficulties in obtaining the model evidence directly, and instead opt for an ad hoc procedure in which a simple heuristic (denoted *L_K_* here) is used as an approximation of –2×log(evidence). The derivation of this statistic rests on certain simplifying assumptions, and the authors are careful to emphasize that these assumptions are ‘dubious’. It is reasonable to expect that any downstream method that relies on the value of *L*_*K*_ (for example Evanno’s Δ*K* statistic (Evanno *et al.* 2005)) is also at the mercy of these dubious assumptions.

Rather than relying on heuristics, what we would like is a direct way of estimating the model evidence that is both accurate and straightforward to implement. As noted by Gelman and Meng (1998), such a method already exists, and has been known about in the physical sciences for some time. This method – referred to in the statistical literature as *thermodynamic integration* (TI) – uses the output of several closely related MCMCs to obtain a direct estimate of the evidence. Crucially, this is not just another heuristic. Rather, it is a true statistical estimator that can be evaluated to an arbitrary degree of precision by simply increasing the number of MCMC iterations used in the calculation. The TI methodology was introduced into population genetics by Lartillot and Philippe (2006) and has since been applied to a range of problems in phylogenetics and coalescent theory (Baele *et al.* 2012; Beerli and Palczewski 2010; Lepage *et al.* 2007; Blanquart and Lartillot 2006), but not yet the problem of estimating *K.*

In the remainder of this paper we demonstrate the effectiveness of TI as a method for estimating *K* in simple genetic mixture models. We find that the Structure estimator *L*_*K*_ is both biased an imprecise, often differing from the true evidence by a factor of 50% or more, while the TI estimator differs by less than 1% for the same computational effort. We also explore the ability of different statistics to correctly estimate *K*, finding that TI outperforms Evanno’s Δ*K*, the Akaike information criterion (AIC), the Bayesian information criterion (BIC) and the Deviance information criterion (DIC). All of the methods described here are made available through the program TrueK (see link at end).

## Methods

### Evidence and Bayes Factors

In a Bayesian setting the problem of deciding between competing models can be addressed using Bayes’ rule. The posterior probability of the model *ℳ*, given the observed data x, can be written

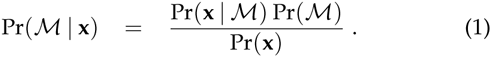

The quantity Pr(**x** | *ℳ*) – the probability of the observed data × given just the model *ℳ* – is defined as the model evidence.

The ratio of the evidence between competing models, known as the *Bayes factor*, can be used to measure the strength of evidence in favour of one model over another. Bayes factors can be used on their own, or they can be combined with priors on the different models to arrive at the posterior odds:

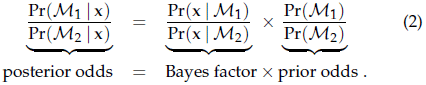

A large Bayes factor in (2) provides evidence in favour of model *ℳ_1_* over model *ℳ_2_*, whereas a small Bayes factor provides evidence in favour of model *ℳ_2_* over model *ℳ_1_*. A useful scale for interpreting Bayes factors can be found in Kass and Raftery (1995).

The problem of estimating the number of demes in a structured population can be understood within this framework. If we let *ℳ*_*K*_ denote a genetic mixture model in which *K* demes are assumed then the problem of estimating the true value of *K* becomes one of comparing between different models. Ideally we would like to solve this problem using the exact model evidence, Pr(**x** | *ℳ*_*K*_). Unfortunately, however, calculating the model evidence in complex, multi-dimensional models is not straightforward. Most of the time we cannot write down the probability of the data under the model without also conditioning on certain known parameters, denoted ***θ***. Obtaining the evidence from the likelihood requires that we integrate over a prior on ***θ***:

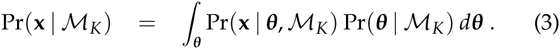

It is this integration step that makes calculating the model evidence difficult in practice. For example, in genetic mixture models ***θ*** might represent the allele frequencies in all *K* demes, perhaps alongside some additional admixture parameters, making the integral in (3) extremely high dimensional (a 100-dimensional integral would not be uncommon). For this reason it is common to turn to numerical methods or heuristic approximations.

### Estimating and Approximating the Evidence

Perhaps the simpest way of estimating the model evidence is through the harmonic mean estimator, 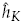 (Newton and Raftery 1994):

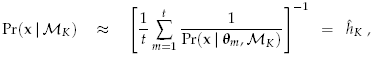

where ***θ***_*m*_ ∈ *m* 6 {1,…, *t*} denotes a series of draws from the posterior distribution of ***θ***. Part of the appeal of this estimator is its simplicity – it is straightforward to calculate 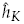 from the output of a single MCMC run. As an example, the program Structurama (Huelsenbeck *et al*. 2011), which contains within it a version of the basic Structure model, has an option for using 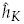 to estimate the model evidence (we note that this is not the primary purpose of Structurama, which also implements a Dirichlet process model). However, in spite of its intuitive appeal, the harmonic mean estimator has been widely criticised to due its instability, 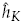 has been found to be very sensitive to the choice of prior, often being dominated by the reciprocal of a few small values (Neal 1994; Raftery *et al.* 2006).

In order to avoid some of the problems inherent in the harmonic mean estimator, the approach taken by Pritchard *et al.* (2000) was to define the heuristic estimator *L_K_* (our notation) as follows:

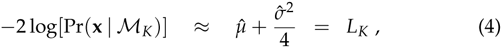

where 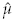 and 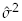 are simple statistics that can be calculated from the posterior draws. The key assumption that underpins this expression is that the posterior deviance is approximately normally distributed (see supplementary text 1 for a more detailed derivation of this and other statistics). *L*_*K*_ can be evaluated for a range of *K*, and the smallest *L*_*K*_ (corresponding to the largest evidence) can be used as an indication of the most likely model. Alternatively, these values can be transformed out of log space to provide direct estimates of the evidence which, once normalised, can be used to approximate the full posterior distribution of *K:*

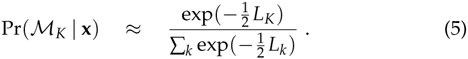

This procedure is rarely carried out in practice, despite being recommended in the Structure software documentation (Pritchard *et al.* 2009).

### Thermodynamic Integration

The TI estimator differs fundamentally from *L*_*K*_ in the sense that it is not a heuristic estimator – it makes no simplifying assumptions about the distribution of the likelihood. It also differs from 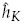 in that it is well-behaved, having finite and quantifiable variance. The approach centres around the ‘power posterior’ (Friel and Pettitt 2008), defined as follows:

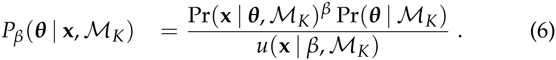

This is nothing more than the ordinary posterior distribution of ***θ***, but with the likelihood raised to the power *β* (the value *u*(**x** | *β*, *ℳ*_*K*_) is a normalising constant that ensures the distribution sums to 1). In the same way that we can design an MCMC algorithm to draw from the posterior distribution of ***θ***, we can design a similar algorithm to draw from the power posterior distribution. Details of the MCMC steps are given in the appendix for models both with and without admixture. The resulting draws from the power posterior will be written 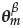, where the superscript *β* indicates the power used when generating the draws. The TI methodology then proceeds in two simple steps. First, we calculate the mean log-likelihood of the power posterior draws:

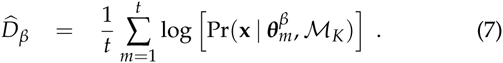

(It is important to note that the notation 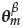, refers to values drawn from the power posterior (with power *β*); it does not indicate that the values of *0* (or these likelihoods) are raised to the power *β*). This step is repeated for a range of values *β_i_*, for *i* = {1,…, *r*} spanning the interval [0,1]. Second, we calculate the area under the curve made by the values 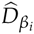 using a suitable numerical integration scheme, such as the trapezium rule:

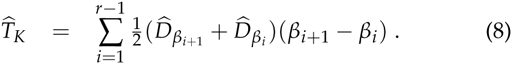

The value 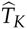 is the TI estimator of the model evidence. It can be seen that 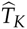 is straightforward to calculate, although it does require us to perform multiple MCMCs to obtain a single estimate of the evidence, making it computationally intensive. On the other hand, the method has greater precision than some alternatives that can be calculated faster. In our comparisons, this trade-off was taken into account by using the same number of MCMC iterations for all methods.

## Results

### Comparison against the exact model evidence

Our first objective was to measure the accuracy and precision of different estimators of the model evidence against the exact value, obtained by brute force (see appendix). The difficulty in calculating the exact model evidence meant that this was only possible for very small simulated data sets of *n* = 10 diploid individuals at *L* = 5 loci, generated from the same without-admixture model implemented in the program Structure2.3.4. A total of 1000 simulated data sets were produced, with *K* ranging from 1 to 10 (100 simulations each). Each data set was then analysed using the program TrueK1.0. This program is written in C++ and was designed specifically to carry out the MCMC procedures described in the appendix. The output of TrueK1.0 includes values of 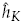, *L*_*k*_, and the TI estimator 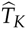 (values of *L*_*K*_ were compared extensively against the output of Structure2.3.4 to ensure agreement). A total of 1000 iterations were obtained from the posterior distribution, with a burn-in of 100 iterations and thinning of 5 iterations. These values are smaller than typical settings for an analysis using the program Structure2.3.4, the reason being that TRUEK1.0 differs in the way it implements the core algorithm, resulting in fewer iterations being needed to ensure approximately independent draws from the posterior (see supplementary text 2 for details). In spite of the apparently small number of samples and burn-in iterations, an autocorrelation analysis indicated that this level of burn-in and thinning was sufficient to ensure pseudo-independent draws from the posterior. For the TI estimator the number of ‘rungs’ used (the number of powers *β*_*i*_,) was set to 50. For 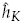 and *L*_*K*_ the analysis was repeated 50 times to obtain a global mean and standard error, thereby ensuring that the same computational effort was expended for all methods.

Figure 1 shows the results of one such analysis, in which the true number of demes was *K* = 2. It can be seen that the Structure estimator *L*_*k*_ is negatively biased in this case, leading to estimates of –2×log(evidence) that are smaller than the true value. This bias is still present after transforming to linear space and normalising results to sum to unity.

**Figure 1.**
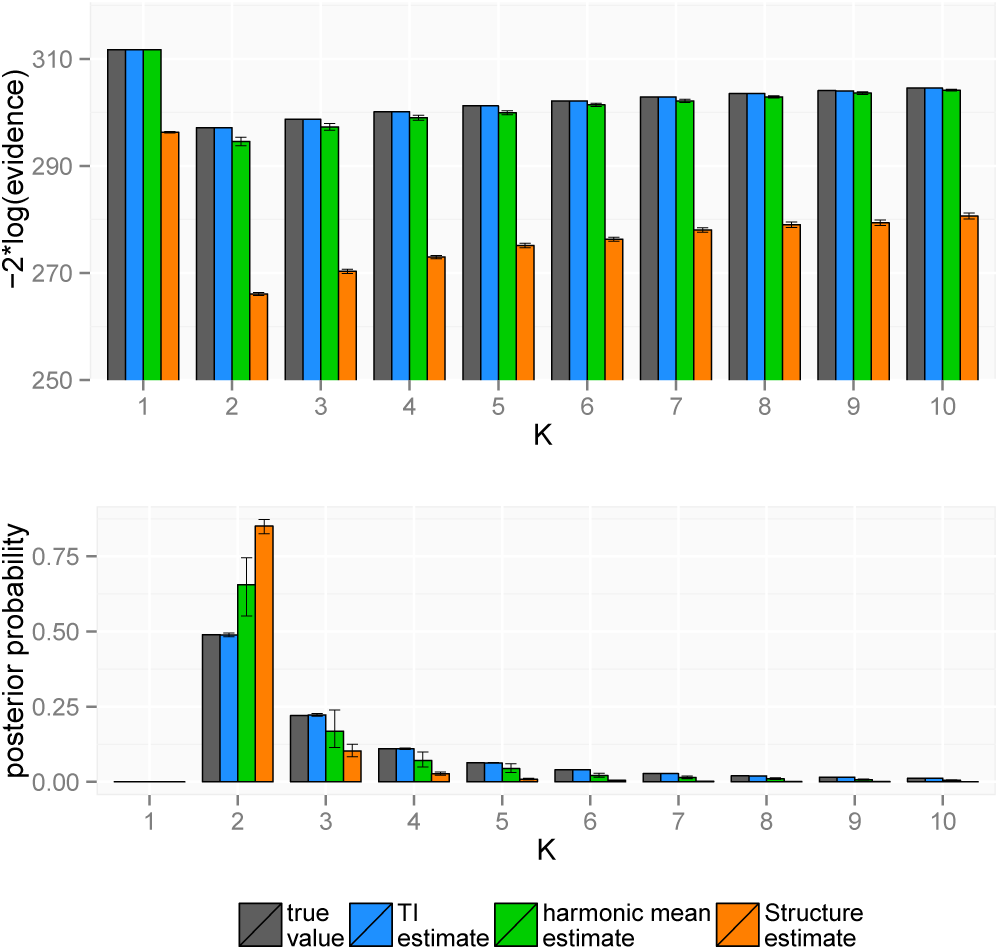
True and estimated values of the model evidence in log space and in linear space. Error bars give 95% confidence intervals around estimates.

The accuracy and precision of the different estimators was evaluated across all 1000 simulated data sets in the form of the Mean Percentage Error (MPE) and the Mean Absolute Percentage Error (MAPE). The MPE measures the average percentage difference between the true and estimated values, and hence can be considered a measure of bias, while the MAPE measures the average absolute percentage difference, and hence can be considered a measure of precision (small values represent more precise estimates). Results are given in Table 1 broken down by the value of *K* used in the model (a more detailed breakdown can be found in supplementary table 3).

**Table 1.** Accuracy of estimation methods compared with the exact model evidence. Values show MPE (MAPE). Normalised values are obtained by first transforming estimates out of log space and then normalising to sum to 1. Values of *K* here denote the value used in the inference step (i.e. the *K* assumed by the model), although multiple values of *K* were used when generating data. A more detailed breakdown can be found in supplementary table 3.

| $K$ | -2log(evidence) | | | normalised evidence | | |
| --- | --- | --- | --- | --- | --- | --- |
| | $\hat{T}_K$ | $\hat{h}_K$ | $L_K$ | $\hat{T}_K$ | $\hat{h}_K$ | $L_K$ |
| 1 | 0.00 (0.00) | 0.00 (0.00) | -4.59 (4.59) | 0.00 (0.35) | -16.34 (16.34) | -93.60 (93.60) |
| 2 | 0.00 (0.01) | -0.19 (0.20) | -7.18 (7.18) | 0.02 (0.95) | 9.89 (14.34) | 70.76 (95.41) |
| 3 | 0.00 (0.01) | -0.23 (0.23) | -7.17 (7.17) | 0.00 (1.00) | 14.30 (16.21) | 38.13 (61.55) |
| 4 | 0.00 (0.01) | -0.19 (0.19) | -7.02 (7.02) | -0.07 (1.01) | 7.36 (10.94) | 1.92 (36.57) |
| 5 | 0.00 (0.01) | -0.15 (0.15) | -6.99 (6.99) | 0.06 (0.94) | 1.53 (8.10) | 0.20 (37.17) |
| 6 | 0.00 (0.01) | -0.12 (0.12) | -7.01 (7.01) | 0.00 (0.89) | -2.62 (7.72) | 6.45 (41.13) |
| 7 | 0.00 (0.01) | -0.10 (0.10) | -7.03 (7.03) | 0.01 (0.88) | -5.71 (8.35) | 15.07 (45.34) |
| 8 | 0.00 (0.01) | -0.07 (0.08) | -7.04 (7.04) | -0.05 (0.86) | -8.24 (9.67) | 22.20 (50.90) |
| 9 | 0.00 (0.01) | -0.07 (0.07) | -7.05 (7.05) | 0.02 (0.79) | -9.22 (10.31) | 27.33 (56.03) |
| 10 | 0.00 (0.01) | -0.05 (0.06) | -7.05 (7.05) | -0.05 (0.83) | -10.65 (11.46) | 31.23 (60.69) |
| mean | 0.00 (0.01) | -0.12 (0.09) | -6.81 (6.81) | -0.01 (0.85) | -1.97 (11.34) | 11.97 (57.84) |

It can be seen that the average MAPE of the *L*_*K*_ estimator across all simulations is 57.84%, while the MAPE of the 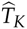 estimator is just 0.85% for the same computational effort. In other words, given an estimate of the model evidence obtained using *L*_*K*_ we can expect the true value to be 57.84% larger or smaller than our estimate on average. The harmonic mean estimator is intermediate between these values, differing from the true evidence by 11.34% on average. Based on these results we would expect estimates of the posterior distribution of *K* made using *L*_*K*_ to be qualitatively different from the true posterior distribution.

### Choosing the ‘best’ *K*

Although the results in Table 1 are suggestive of a weakness in the *L*_*K*_ estimator, we are limited here to looking at small data sets in which the exact model evidence can be calculated by brute force. It is plausible based on these results that the bias in *L*_*K*_ could be amplified in small data sets due to a lack of information, and would cease to be a problem if more data were available. There is also a question of whether it matters that *L*_*K*_ fails to estimate the model evidence as long as the single ‘best’ value of *K* is correctly identified. For example, in Figure 1 all estimation methods correctly identify that *K =* 2 is the most likely value, even though our confidence in that result may be misplaced in some cases.

Here we therefore use larger simulated data sets to address the question of whether the TI method produces improvements that would be of practical importance. To begin with, we compare the ability of different methods to estimate the single ‘best’ value of *K*, irrespective of the logic behind the statistic. By using simulated data sets, in which a known value of *K* is used when generating the data, we can measure the proportion of times that the true value is correctly identified. As well as comparing the estimators 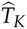, 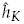 and *L*_*K*_, in which the smallest value of the estimator indicates the most likely model, we also compare Evanno’s Δ_*K*_ (Evanno *et al.* 2005), in which the largest value indicates the point of maximum curvature of *L*_*K*_, and the AIC, BIC and DIC statistics, in which the smallest value indicates the best fitting model. Values of the DIC are calculated using the method of Spiegelhalter *et al.* (2002) (DIC_S_) as well as the method of Gelman *et al.* (2014) (DIC_G_). To ensure that our results are not driven by a lack of information, larger data sets of *n =* 100 diploid individuals at *L* = 10 loci were generated from the same without-admixture model used above. As before, 1000 simulated data sets were produced, with *K* ranging from 1 to 10 (100 simulations each). TrueK1.0 was run with 200 burn-in iterations, 1000 sampling iterations and thinning every 5 iterations. As before, an autocorrelation analysis indicated that these parameter values were sufficient to ensure pseudo-independent draws from the posterior. For the TI estimator 50 rungs were used, and for *L*_*K*_ and 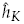 the analysis was repeated 50 times.

Table 2 gives the number of times that the correct value of *K* was identified by each of the methods. As there are 100 simulations for each *K*, these values are also equivalent to percentages. It can be seen that the TI based method of choosing *K* provides the most reliable results, with the majority of simulated data sets being correctly identified over a wide range of *K.* Estimates of *K* based on 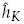 are less reliable, leading to incorrect conclusions in a greater proportion of simulations, and estimates based on *L*_*K*_ are even more unreliable, with a clear bias towards large values of *K.* Evanno’s Δ_*K*_ shows the opposite trend, favouring small values of *K* (although Δ_*K*_ can never predict *K* = 1). Of the model comparison statistics the DIC calculated using the method of Gelman *et al.* (2014) (DIC_G_) appears to be the most reliable, although this is still outperformed by TI. Surprisingly the BIC, which has been found to give reliable results in other mixture modelling applications (Steele and Raftery 2009; Fraley and Raftery 1998), performs poorly here.

**Table 2.** Number (%) of times K correctly identified.

| $K$ | $\hat{T}_K$ | $\hat{h}_K$ | $L_K$ | $\Delta_K$ | AIC | BIC | DIC<sub>s</sub> | DIC<sub>G</sub> |
| --- | --- | --- | --- | --- | --- | --- | --- | --- |
| 1 | 100 | 92 | 0 | 0 | 85 | 100 | 0 | 97 |
| 2 | 100 | 91 | 0 | 100 | 97 | 100 | 0 | 88 |
| 3 | 100 | 84 | 2 | 73 | 98 | 99 | 0 | 87 |
| 4 | 100 | 64 | 15 | 59 | 97 | 77 | 0 | 80 |
| 5 | 100 | 66 | 21 | 47 | 100 | 26 | 0 | 89 |
| 6 | 96 | 64 | 31 | 24 | 95 | 3 | 2 | 83 |
| 7 | 91 | 57 | 35 | 12 | 66 | 0 | 2 | 82 |
| 8 | 73 | 52 | 40 | 4 | 28 | 0 | 8 | 64 |
| 9 | 60 | 47 | 46 | 0 | 4 | 0 | 25 | 49 |
| 10 | 89 | 91 | 88 | 0 | 0 | 0 | 93 | 40 |

Returning to the question of whether the inaccuracy in 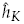 and *L*_*K*_ in Table 1 was driven by a lack of information, it can be seen from Table 2 that even for reasonably large data sets the bias and error in the 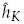 and *L*_*K*_ statistics can lead to qualitatively different conclusions about *K.* Thus, the increased precision of the TI approach is of practical as well as theoretical importance.

### Comparing between different evolutionary models

As well as being useful for inferring *K*, the model evidence provides a robust framework for comparing between different evolutionary models. The analyses described so far have all involved a model comparison of sorts, in the sense that each *K* technically represents a different model *ℳ*_*K*_, but the evolutionary model (i.e. the without-admixture model) has been assumed known. However the evolutionary model will often be of greater interest; for example, a biologist might be less interested in the exact number of subpopulations, but rather in the level of genetic admixture between them. In such circumstances, *K* can be thought of as indexing a series of models within a wider evolutionary model, *ℳ.* We can compare models at this wider level by simply integrating *K* out as a nuisance factor:

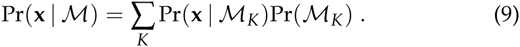

If we assume an equal prior over *K* then this amounts to taking the mean of the evidence for each value of *K* to arrive at the overall evidence for *ℳ.*

To explore this point, a single simulated data set of *n =* 100 diploid individuals at *L* = 5 loci (i.e. relatively weak information per individual) was generated from the without-admixture model with *K* = 5 demes. This data set was then analysed under three competing models; the without-admixture model (i.e. the correct model), the with-admixture model used by Structure2.3.4 with admixture parameter α = 1.0, and the with-admixture model used by Structure2.3.4 where *α* is inferred as part of the MCMC (given a Uniform(0,10) prior). All three models were run using the program TRUEK1.0. In all cases burn-in and thinning values were chosen based on an autocorrelation analysis, leading to burn-in={200,500,5000} and thinning={5,10,100} for the three models respectively, and in all cases 10,000 draws were obtained from the posterior distribution.

The results of this analysis are shown in Figure 2 for each of the three evolutionary models. Looking first at the overall evidence for each evolutionary model (Figure 2 E) we can see that the without-admixture model obtains the highest overall evidence, followed by the admixture model with *α* free to vary. We can dig deeper into the latter model by looking at the posterior distribution of α (Figure 2 D). Very small values of *α* are observed for *K* = {3,4,5}, which are the most likely values of *K* under this model (as shown in Figure 2 C). When *α =* 0 this model becomes directly equivalent to the without-admixture model, and so the small values of *α* here indicate that admixture is very weak, if it is present at all (in fact we know that it is not). Finally, looking at the posterior distribution of *K* in the without-admixture model (Figure 2 A) we can see that *K* = {4,5,6} are all plausible within this model. Note that although the Structure estimator *L*_*K*_ estimates the correct value of *K =* 5 under the without-admixture model, in general neither *L*_*K*_ nor 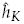 can be relied upon to give reliable results in this example. Thus, using current methods it is not clear whether admixture is small or zero, and even then the value of *K* may be estimated incorrectly, whereas using TI we can make a direct comparison between evolutionary models and find convincing evidence for the correct model.

**Figure 2.**
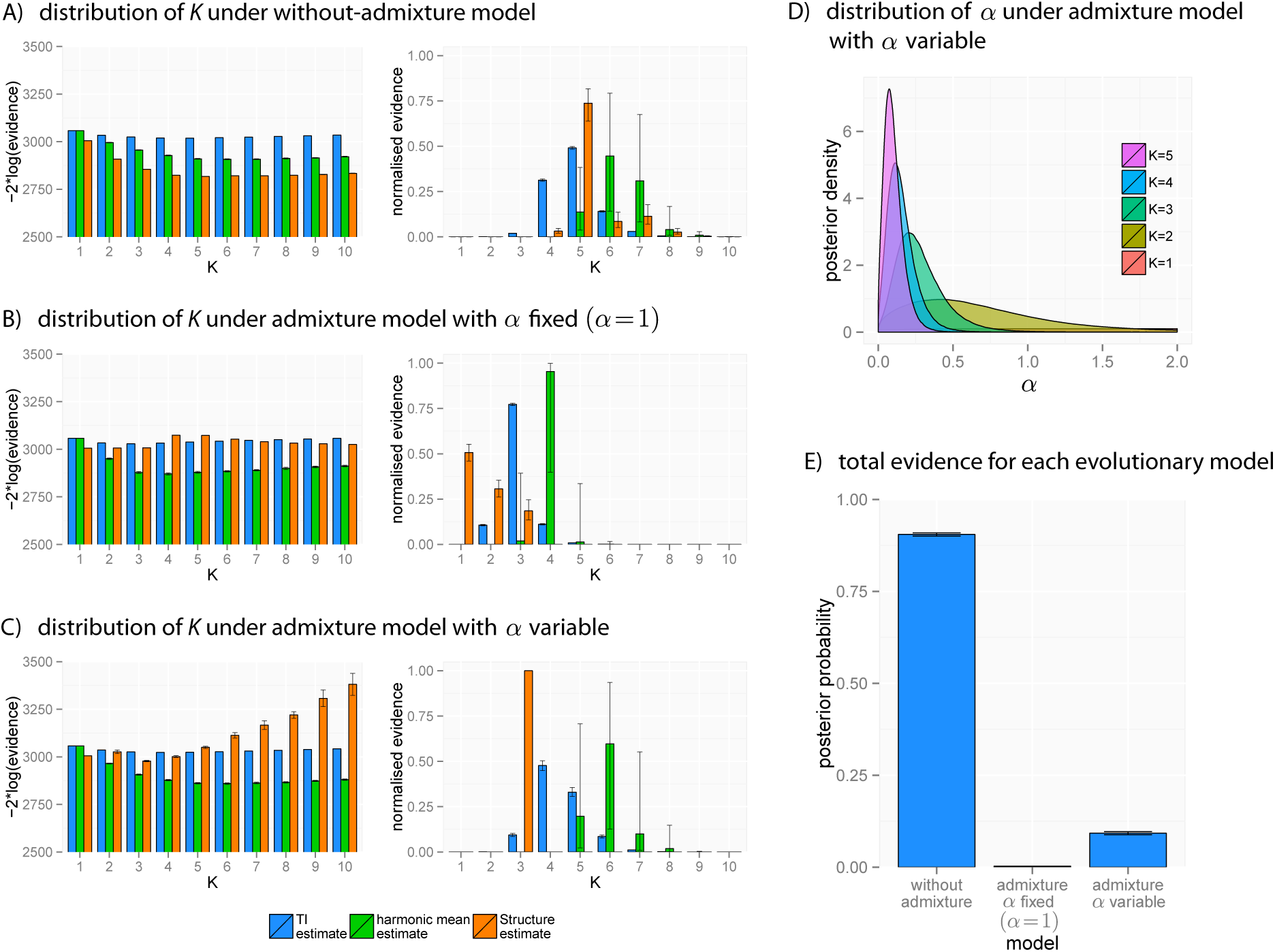
Analysis of simulated data under three evolutionary models. A), B), C) evidence for *K* under all three evolutionary models in log space and linear space after normalisation, D) posterior distribution of *α* under the variable it admixture model (truncated at *α* = 2.0, although distributions extend to α = 10.0), E) overall evidence for each evolutionary model, obtained by summing the evidence for *K* = {1,…, 10} given an equal Pr(*K*) = 0.1 prior for each model.

## Discussion

Model based clustering methods have proved extremely useful within population genetics. The probabilistic allocation of individuals to demes employed by programs such as Structure has made it possible to tease apart population subdivision within a wide range of organisms, including humans (Rosenberg *et al.* 2002; Li *et al.* 2008; Tishkoff *et al.* 2009), human pathogens (Falush *et al.* 2003b), plants (Garris *et al.* 2005) and animals (Parker *et al.* 2004). However, these posterior assignments are always produced conditional on the known value of *K.* Choosing an appropriate value of *K* is statistically much more challenging than estimating population assignments, as it involves a comparison between models rather than simple parameter estimation within a given model. A standard way of comparing models within a Bayesian setting is through the model evidence – defined as the likelihood of the model integrated over all free parameters. Indeed, the Structure estimator *L*_*K*_ was originally designed as an approximation to the model evidence, although we show above that this approximation can be quite poor in some cases. Thermodynamic integration offers a way out of this problem, providing estimates of the model evidence that are both accurate and precise (Table 1), and in turn leading to estimates of *K* that are more reliable than other methods (Table 2).

Although the results above are promising, when thinking about population structure it is important that we do not place too much emphasis on any single value of *K.* The simple models used by programs such as Structure and TRUEK1.0 are nothing more than highly idealised cartoons of real life, and so we cannot expect the results of model-based inference to be a perfect reflection of true population structure (see discussion in Waples and Gaggiotti (2006)). Thus, while TI can help ensure that our results are statistically valid conditional on a particular evolutionary model, it is still possible to generate results that are biologically meaningless if the evolutionary model is not appropriate for the data. Similarly – in spite of the results in Table 2 – we do not advocate using the model evidence (estimated by TI or any other method) as a way of choosing the single ‘best’ value of *K.* The chief advantage of the evidence in this context is that it can be used to obtain the complete posterior distribution of *K*, which is far more informative than a single point estimate. Although one value of *K* may be most likely *a posteriori*, in general a range of values will be plausible, and we should entertain all of these possibilities when drawing conclusions. As a concrete example, even for those simulations in Table 2 where TI failed to estimate the correct *K*, the true value of *K* was within the 95% credible interval 94.5% of the time.

Looking at the wider issue of which evolutionary model is most appropriate for our data, once again the model evidence can be an extremely useful tool here. By treating *K* as a nuisance factor, choosing to integrate it out of the problem rather than focussing on it, we can begin to compare models at this wider level. In doing so we guard against the temptation to apply a single evolutionary model to all data types, irrespective of whether the model is in fact appropriate for the data at hand. This is particularly important when our focus is on inferring the presence or absence of admixture between subpopulations, as the particular form of the model may play a large role in whether or not we detect a signal of admixture.

In either case – whether we are looking at the distribution of *K* within an evolutionary model, or integrating over *K* as a nuisance factor – the ability to obtain reliable estimates of the model evidence is a prerequisite, and TI takes us one more step in this direction.

The TRUEK1.0 program and documentation can be downloaded from www.bobverity.com/TrueK

## Appendix

### MCMC under the without-admixture model

In order to carry out the TI estimation approach we need to be able to draw from the power posterior distribution. This is straightforward in the case of genetic mixtures, and requires nothing more than a simple extension of existing MCMC algorithms. In the following we strive to bring our notation in line with previous studies wherever possible, but the complexities of certain likelihood functions also motivate us to define some new notation (see Table 3). It is worth noting, for example, that we will write individual genotypes in simple list form (as in Pritchard *et al.* (2000)) using the notation *x_il_* for the *l*^th^ locus of the *i*^th^ individual, but also in allelic partition form (as in Huelsenbeck *et al.* (2011)) using the notation ***s***_*il*_. For example, a diploid individual homozygous for the third allele at a particular locus can be written ***x***_*il*_ = (3,3) or equivalently ***s***_*il*_ = {0,0,2,0,0}, where there are five possible alleles to choose from in this example. Conditioning on the model ℳ_*K*_ is also implicit throughout this section. In the basic algorithm described by Pritchard *et al.* (2000) there are two free parameters to keep track of – the allocation of individuals to demes, denoted **z** here, and the allele frequencies in all *K* demes, denoted **p.** Under the assumptions of Hardy-Weinberg and linkage equilibrium it is possible to write down the probability of the observed data given the known values of these free parameters, Pr(**x** | **z**, **p**). Combining this likelihood with a Dirichlet(λ_1_… λ_*J_l_*_) prior on the allele frequencies at each locus we can derive the conditional posterior distribution of the allele frequencies given the known group allocation, Pr(**p** | **x**, **z**). Alternatively, multiplying by an equal *1/K* prior on the allocation of individuals to demes we can derive the conditional posterior distribution of the group allocation given the known allele frequencies, Pr(**z** | **x**, **p**). Algorithm 1 of Pritchard *et al.* (2000) works by alternately sampling from each of these conditional distributions, resulting (after sufficient burn-in) in a series of draws from the full posterior distribution. More often than not we are interested in the posterior allocation, in which case the posterior allele frequencies can simply be ignored.

**Table 3.** Definitions of parameters used in this study.

| parameter | description |
| --- | --- |
| $K$ | No. of populations |
| $L$ | No. of loci |
| $J_l$ | No. of unique alleles observed at locus $l$ |
| $n$ | No. of individuals sampled |
| $\mathbf{x}$ | Allele information for all individuals |
| $\mathbf{x}_i$ | Allele information for individual $i$ |
| $\mathbf{x}_{il}$ | Allele information for individual $i$ at locus $l$ |
| $x_{ila}$ | Allelic type of the $a^{\text{th}}$ gene copy in individual $i$ at locus $l$<br>[$x_{ila} \in (1, \dots, J_l)$] |
| $\mathbf{s}_{il}$ | Allelic partition of alleles in individual $i$ at locus $l$. For example, the genotype $\mathbf{x}_{il} = (3, 3)$ can be written $\mathbf{s}_{il} = \{0, 0, 2, 0, 0\}$ in allelic partition form, where in this example $J_l = 5$ |
| $s_{ilj}$ | No. of copies of allele $j$ at locus $l$ in individual $i$ |
| $s_{il}$ | No. of copies of any allele at locus $l$ in individual $i$ |
| $\mathbf{y}_k$ | Allelic partition at all loci of all gene copies assigned to population $k$ |
| $\mathbf{y}_{kl}$ | Allelic partition at locus $l$ of all gene copies assigned to population $k$ |
| $y_{klj}$ | No. of copies of allele $j$ at locus $l$ assigned to population $k$ |
| $y_{kl}$ | No. of copies of any allele at locus $l$ assigned to population $k$ |
| $\mathbf{p}$ | Allele frequencies in all populations at all loci |
| $z_{ila}$ | Assignment of gene copy $a$ at locus $l$ in individual $i$ to a population [$z_{ila} \in (1, \dots, K)$] |
| $z_i$ | Assignment of individual $i$ to a population. When referring to $z_i$ it is implied that $z_{ila}$ is identical for all $(l, a)$, meaning all gene copies within this individual are assigned together |
| $\mathbf{z}$ | Assignment of all gene copies in all individuals |
| $\mathbf{q}_i$ | Admixture proportions in individual $i$ |
| $\mathbf{v}$ | Partition of gene copies to populations in all individuals |
| $\mathbf{v}_i$ | Partition of gene copies to populations in individual $i$ |
| $v_{ik}$ | No. of gene copies in individual $i$ assigned to population $k$ |
| $v_i$ | No. of gene copies in individual $i$ assigned to any population |
| $\lambda_{lj}$ | Dirichlet parameter for frequency of allele $j$ at locus $l$ |
| $\lambda_{l0}$ | Sum of the Dirichlet parameters for locus $l$ [$\lambda_{l0} = \sum_{j=1}^{J_l} \lambda_{lj}$] |
| $\alpha$ | Dirichlet parameter on admixture proportions |
| $c^{(-i)}$ | Evaluation of a parameter while excluding information for individual $i$ ($c$ could be any of the parameters listed above) |
| $c^{(-ilp)}$ | Evaluation of a parameter while excluding information for gene copy $a$ at locus $l$ in individual $i$ ($c$ could be any of the parameters listed above) |
| $\Gamma(\cdot)$ | Gamma function |

However, as stated in the original derivation of Rannala and Mountain (1997) and reiterated by Huelsenbeck *et al.* (2011), it is possible to integrate over the allele frequencies analytically, thereby greatly reducing the dimensionality of the problem. The new likelihood, conditional only on the group allocation, can be written

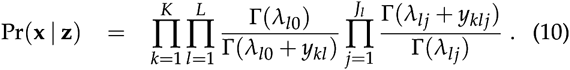

This expression is extremely useful to us, as it means the likelihood can be calculated without having to take into account an explicit representation of the unknown allele frequencies – our uncertainty in the allele frequencies has already been integrated out of the problem.

Rather than using (10) directly, Huelsenbeck *et al.* (2011) used this analytical solution to define an efficient MCMC algorithm. Dividing the probability of the data **x** by the probability of the data with the *i*^th^ observation removed, denoted ***x***^(*-i*)^ we obtain the conditional probability of observation *i* given all others. This can be written

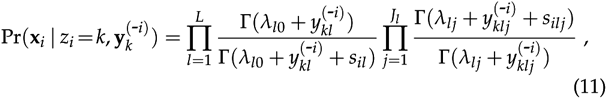

which is equivalent to formula 4 in Huelsenbeck *et al.* (2011), although derived via a slightly different route. Computing (11) for all *k* and normalising we obtain the conditional posterior probability that individual *i* belongs to deme *k:*

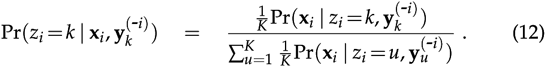

By repeatedly drawing new group allocations for all individuals from (12) we obtain a series of draws from the posterior distribution without ever needing to invoke the unknown allele frequencies. Thus, the two-step algorithm of Pritchard et al. Pritchard *et al.* (2000) can be reduced to the more efficient one-step algorithm of Huelsenbeck *et al.* (2011).

We can make use of the same gains in efficiency when designing an MCMC algorithm for the purposes of TI. In fact, the only difference when carrying out TI is that the likelihood in (10) should be raised to the power *β*, allowing us to draw from the power posterior. On making this change we find that the conditional posterior distribution in (11) should also be raised to the power *β* (this follows from the fact that (11) can be derived as a ratio of two ordinary likelihoods). Thus, we arrive at a new expression for the probability of individual *i* being assigned to group *k:*

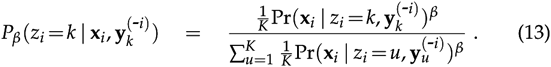

By repeatedly sampling new group allocations for all individuals from (13) we obtain a series of allocation vectors drawn from the power posterior (notice that when *β* = 0 we are essentially drawing from the prior). The likelihood of each vector can then be computed using (10), at which point we have everything we need to calculate 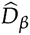 as in (7). Carrying out this entire procedure for a range of values *β*_*i*_, we obtain a series of points *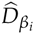.* Which can be used to calculate the TI estimator 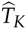, as in (8). The complete TI algorithm for the model without admixture can be defined as follows:

#### Algorithm 1 (without admixture)

1. For *r* distinct values of *β*_*i*_, spanning the interval [0,1]
  a. Perform MCMC by repeatedly drawing from (13) for all *i* ∈ {1,…,*n*} This results (after discarding burn-in and thinning) in *t* approximately independent draws from the power posterior group allocation.
  b. Calculate the likelihood of each group allocation using (10).
  c. Calculate *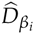.* as the average log-likelihood, as in (7).
2. Use the values *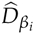.* to calculate 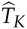 in a suitable numerical integration scheme, for example using the trapezium rule as in (8).

### MCMC under the admixture model

The model with admixture described by Pritchard *et at.* (2000) is slightly complicated by the fact that each gene copy is free to originate from a different deme. However, we can still apply the same basic logic described above to arrive at a simple one-step algorithm for sampling from the power posterior. First we note that the probability of the data conditional on the known group allocation is identical in this model to the probability in the without-admixture model, and is given by (10). This is true because we make the same assumption of gene copies being drawn independently from demes, and we apply the same Dirichlet priors on allele frequencies, meaning the final likelihood does not change. The difference in the admixture model is that the group allocation takes place at the level of the gene copy, rather than at the level of the individual, and so the values *Z*_*ila*_ are no longer restricted to being the same for all (*l*, *a*). This is reflected in the **y**_*k*_ values used to keep track of the gene copies allocated to a particular deme, which are now free to contain only a partial contribution of the genome of each individual.

Following the same approach as for the without-admixture model, we can obtain the conditional probability of gene copy *x*_*ila*_ by dividing through the probability of the complete data set by the probability of the data set with this element removed (denoted ***x***^(*–ila*)^). Most of the terms in the resulting expression cancel out, leading to the following simple result:

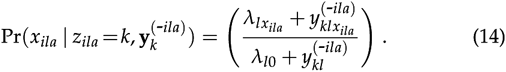

As before, this likelihood should be combined with the prior probability of assignment to each deme. If the admixture proportions for individual *i* are given by the vector **q**_*i*_, then, under the assumptions of the model described by Pritchard *et al.* (2000), the number of gene copies in this individual that are allocated to each deme can be considered a multinomial draw from **q**_*i*_,. Integrating over a Dirichlet(*α*, *α*,…*α*) prior on these frequencies we obtain

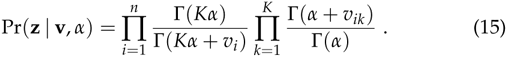

We can use this expression to write down the prior probability of a single gene copy being allocated to deme *k:*

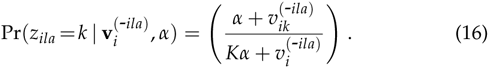

Bringing together the prior with the likelihood raised to the power *β* we obtain the following expression for the power posterior probability of an individual gene copy being allocated to deme *k*:

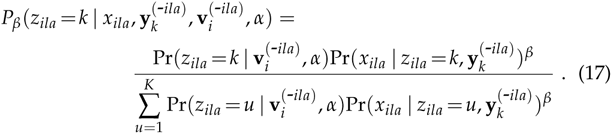

By repeatedly sampling new allocations for all gene copies at all loci within all individuals (i.e. all *z*_*ila*_) we obtain a series of draws from the power posterior group allocation under the admixture model. Again, this algorithm is made more efficient by the fact that the unknown allele frequencies in all populations and the unknown admixture proportions in all individuals have been integrated out of the problem at an early stage.

A common extension to the basic admixture model is to leave α as a free parameter, updating it as part of the MCMC. This can be accommodated within the TI framework by using a simple Metropolis-Hastings step. If *α’* is a new value of *α*, drawn from some suitable proposal distribution *g*(*α*’ | *α*), then the acceptance probability under Metropolis-Hastings is given by

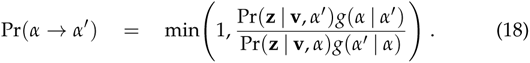

Notice that the core probability that drives this expression is the *prior* probability of the allocation **z**, which is given in (15). The actual probability of the data – i.e. the expression that is raised to the power in the power *β* posterior calculation – does not feature here. Thus, we can use the same Metropolis-Hastings step to update α irrespective of the value of *β*.

The complete TI algorithm for the model with admixture can be defined as follows:

#### Algorithm 2 (with admixture)

1. For *r* distinct values of *β*_*i*_, spanning the interval [0,1]
  (a) Perform MCMC by repeatedly drawing from (17) for all gene copies at all loci in all individuals (all *a, l, i*). If *α* is a free parameter then update this value using a Metropolis-Hastings step, as in (18). This results (after discarding burn-in and thinning) in ***t*** approximately independent draws from the power posterior group allocation.
  (b) Calculate the likelihood of each group allocation using (10).
  (c) Calculate *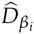.* as the average log-likelihood, as in (7).
2. Use all the values 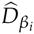 to calculate 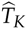 in a suitable numerical integration scheme, for example using the trapezium rule as in (8).

Finally, we note that the expressions derived in this section can be used to obtain the exact model evidence by brute force in restricted settings. For example, focussing on the model without admixture, we could sum over the likelihood of all possible group allocations to obtain the true model evidence:

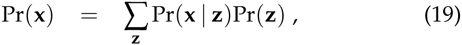

where Pr(**x** | **z**) is given by (10), and for this model Pr(**z**) = 1/*K^n^* for all group allocations. Although this is possible in theory, the sheer number of allocations that we are required to sum over makes this method impractical in all but the simplest situations. Even if we exploit redundancies in the labelling of different allocations we are still restricted to values of *n* and *K* not much larger than 10. This method is therefore only really useful as a way of checking the accuracy of other estimation methods.

